# Integrated genomics provides insights for the evolution of the polyphosphate accumulation trait of *Ca.* Accumulibacter

**DOI:** 10.1101/2023.09.20.558572

**Authors:** Xiaojing Xie, Xuhan Deng, Liping Chen, Jing Yuan, Hang Chen, Chaohai Wei, Xianghui Liu, Stefan Wuertz, Guanglei Qiu

## Abstract

*Candidatus* Accumulibacter plays a major role in enhanced biological phosphorus removal (EBPR), but the key genomic elements in metagenome assembled genomes enabling their phosphorus cycling ability remain unclear. Pangenome analyses were performed to systematically compare the genomic makeup of *Ca.* Accumulibacter and non-*Ca*. Accumulibacter members within the Rhodocyclaceae family. Metatranscriptomic analyses of an enrichment culture of *Ca.* Accumulibacter clade IIC strain SCUT-2 were performed to investigate gene transcription characteristics in a typical anaerobic-aerobic cycle. Two hundred ninety-eight core genes were shown to be obtained by *Ca.* Accumulibacter at their least common ancestor. One hundred twenty-four of them were acquired via horizontal gene transfer (HGT) based on best-match analysis against the NCBI database. Fourty-four laterally derived genes were actively transcribed in a typical EBPR cycle, including the polyphosphate kinase 2 (PPK2) gene. Genes in the phosphate regulon (Pho) were poorly transcribed. Via a systematical analysis of the occurrences of these genes in closely related *Dechloromonas*-polyphosphate accumulating organisms (PAOs) and *Propionivibrio*-non-PAOs, a Pho dysregulation hypothesis is proposed to explain the mechanism of EBPR. It states that the PhoU acquired by HGT fails in regulating the high-affinity phosphate transport (Pst) system. To avoid phosphate poisoning, the laterally acquired PPK2 is employed to condense excess phosphate into polyphosphate. Alternatively, genes encoding PhoU and PPK2 are obtained from different donor bacteria, leading to unmatched phosphate concentration thresholds for their activation/inactivation. PPK2 tends to reduce the intracellular phosphate to concentration levels perceived by PhoU as low-phosphate states. PhoU is not activated to turn off the Pst system, resulting in continuous phosphate uptake. In conclusion, based on integrated genomic analyses, the HGT of *pho*U and *ppk*2 and the resultant Pho dysregulation may have triggered the development and evolution of the P cycling trait in *Ca.* Accumulibacter.

## 1 Introduction

With the rapid development of industry and the economy, increased amounts of wastewater are generated. The resultant excess discharge of phosphorus (P) leads to eutrophication, deteriorated water quality, and aquatic ecosystem degeneration (Bunce et al., 2018; Abdelfattah et al.,2023; Qiu et al., 2022; Liu et al., 2023). Enhanced biological phosphorus removal (EBPR) is an environmental-friendly and economical process widely applied in municipal wastewater treatment plants (WWTPs) for P removal (García Martín et al., 2006; Oehmen et al., 2007; Qiu et al., 2019; 2020; Wang et al., 2021; Chen et al., 2022; Diaz et al., 2022; Zhang et al., 2022). This process is mediated by a group of microorganisms namely polyphosphate-accumulating organisms (PAOs) (He and McMahon, 2011; Nielsen et al., 2019; Dorofeev et al., 2020; Zhang et al., 2023;). *Candidatus* Accumulibacter is a model genus of PAOs commonly found in lab- and full-scale EBPR systems (Seviour et al., 2003; Mao et al., 2015; Roy et al., 2021; Petriglieri et al., 2022; Deng et al., 2023). Under anaerobic conditions, *Ca.* Accumulibacter use the intracellularly stored polyphosphate (poly-P) as an energy source to power the uptake of volatile fatty acids (VFAs). Phosphate is released. The assimilated VFAs are polymerized and stored as polyhydroxyalkanoates (PHAs). In the subsequent aerobic phase, PHAs are oxidized for cell metabolism and reproduction. Excess phosphate is taken up from the aquatic phase to synthesize poly-P, achieving P removal (Oehmen et al., 2005; Kolakovic et al., 2021; Zhao et al., 2022; Páez-Watson et al., 2023). This unique metabolic feature allows PAOs to thrive in alternating anaerobic-aerobic conditions, conferring sustainable P removal. However, the key genetic basis affording PAOs the ability of P cycling is not clear. Genes known to be indispensable for the P cycling feature, e.g., the polyphosphate kinase gene (*ppk*) and exopolyphosphatase gene (*ppx*) for poly-P synthesis and hydrolysis, respectively, and the inorganic phosphate transporter gene (*pit*) and the high-affinity phosphate transporter gene (*pst*) for phosphate transport, are widely preserved in the bacterial domain, including in non-PAOs (Bessarab et al., 2022; Maszenan et al., 2022). Their presence does not guarantee the P cycling ability and the key genes have yet to be identified. The transition from non-PAOs to PAOs may be driven by adaptive evolution (Turcotte et al., 2012; Oyserman et al., 2016a).

The need to understand the gain and loss of genes in different strains and the genome diversification in a given lineage of organisms gave rise to the concept of pangenomics. A pangenome is the entire set of genes from all individuals of a specific lineage (Tettelin et al., 2005; Song et al., 2020). Genes in a pangenome are divided into core genes and variable genes (Medini et al., 2005). The collection of genes which are commonly present in all individuals of a specific lineage is called the core pangenome, representing the common genetic features of a microbial lineage (Della et al., 2021). The variable genes can be further divided into unique genes (found in a single strain/genome) and dispensable genes (shared in at least 2 but not all strains/genomes) (Aggarwal et al., 2022). Dispensable genes represent the intra-lineage diversity encoded among different members (Medini et al., 2005). By avoiding single sample bias and ensuring full representation of genomic diversity of different lineage members, the analysis of the pangenome provides insight into the genetic basis of common phenotypic characteristics shared in a group of bacteria, greatly improving our ability to solve complex phenotypic problems (Golicz et al., 2016, Flowers et al., 2013; Pamela et al., 2019). Comparative genomics has been applied to study the evolution and development of many bacterial species (El-Sayed et al., 2005; Fernández-Gómez et al., 2013; Coghlan et al., 2019; Kjærbølling et al., 2020; Feng et al., 2020; Zhang et al., 2022). Via comparative genomic analysis, Fernandez-Fueyo et al. (2012) found a subset of potentially important genes for selective lignin decomposition in *Ceriporipsis subvermispora*.

Oyserman et al. (2016a) previously constructed a pangenome of the Rhodocyclaceae family (including 10 *Ca.* Accumulibacter and 16 out-group genomes) to explore the genetic composition and evolutionary changes in metabolic pathways of the *Ca.* Accumulibacter genus. However, at the time, limited numbers of *Ca.* Accumulibacter genomes were available with more than half of the genomes having low completeness (< 90%). The deficiency in genome quality and quantity may result in inadequate representation of the lineage pangenome and affect downstream analysis of genes. With the advance in EBPR research and sequencing techniques, increasing numbers of high-quality *Ca*. Accumulibacter genomes have been obtained (Arumugam et al., 2019; Rubio-Rincón et al.; 2019; Qiu et al., 2020; McDaniel et al., 2021; Singleton et al., 2021; Srinivasan et al., 2021; Tian et al., 2022; Petriglieri et al., 2022; Zhang et al., 2023; Deng et al, 2023). Additionally, new PAOs and glycogen accumulating organisms (GAOs) were identified in genera phylogenetically closely related to *Ca*. Accumulibacter. GAOs occupy a similar ecological niche as PAOs in EBPR systems.

They use glycogen instead of polyphosphate as an energy source for anaerobic carbon sources uptake, thus competing with PAOs. For instance, a *Propionivibrio* member was shown to perform as a GAO in full-scale WWTPs in Denmark (Albertsen et al., 2016). Two *Dechloromonas* members in the same WWTPs (i.e., *Ca.* Dechloromonas phosphoritropha and *Ca.* Dechloromonas phosphorivorans) were revealed to be PAOs (Petriglieri et al., 2021). The identification of *Dechloromonas*-related PAOs raises the possibility that the emergence of the PAO phenotype may have occurred before the *Ca.* Accumulibacter last common ancestor (LCA). The evolution in the P cycling feature needs to be re-evaluated and traced. Combined with the analysis of gene transcriptional characteristics of representative PAO strains, the key genomic characteristics distinguishing PAOs and non-PAOs may be further identified and determined, which would significantly advance the understanding of the genomic basis of the PAO phenotype.

To understand the emergence of the PAO phenotype of *Ca.* Accumulibacter, we selected 43 high-quality genomes within the Rhodocyclaceae family for comparative genomic analysis. A pangenome of the Rhodocyclaceae family including 21 *Ca*. Accumulibacter genomes, seven of which were recovered from our EBPR reactors, 22 out-group genomes, including two confirmed *Dechloromonas* PAOs, i.e., *Ca.* Dechloromonas phosphoritropha and *Ca.* Dechloromonas phosphorivorans (Petriglieri et al., 2021), and one *Propionivibrio* GAO genome, *Ca.* Propionivibrio aalborgensis (Albertsen et al., 2016), was constructed. Genes in the pangenome were classified as ancestral, derived, flexible, or lineage-specific genes. The dynamics in these genes in the evolutionary process were analyzed and metatranscriptomic analyses were performed on an enrichment culture of *Ca*. Accumulibacter Clade IIC SCUT-2 for the identification of their active genes in a typical anaerobic-aerobic cycle to narrow down the range of genes important for the PAO phenotype of *Ca*. Accumulibacter. Genomic comparisons were further performed between *Ca*. Accumulibacter, two *Dechloromonas*-related PAOs, and the *Propionivibrio* GAO. The phosphate signaling complex protein gene (*pho*U) in the Pho regulon and the laterally derived polyphosphate kinase 2 gene (*ppk*2) were determined to have played a crucial role in the emergence of the PAO phenotype of *Ca.* Accumulibacter. This study provides new insights into the development of the P cycling trait of *Ca.* Accumulibacter.

## 2 Materials and Methods

### 2.1 Data acquisition and evaluation

The genomes used for analysis included 7 high-quality genomes recovered from our EBPR reactors and 36 genomes obtained from the National Center for Biotechnology Information (NCBI) database. All 43 genomes belong to the Rhodocyclaceae family, including 21 *Ca.* Accumulibacter genomes and 22 out-group genomes (10 *Dechloromonas*, 7 *Thauera*, 3 *Azoarcus*, 1 *Propionivibrio* and 1 *Zooglea ramigera* genomes). The completeness and contamination of the genomes were evaluated using CheckM (Parks et al., 2015). The GenBank assembly accession, corresponding species names, and additional details about the qualities of these genomes can be found in the Supplementary Material Table S1-S3.

### 2.2 Orthologue analyses

Orthologous gene clustering is necessary for the reconstruction of the ancestral state. To find orthologous gene clusters based on the protein sequences, all vs all BLAST of each Rhodocyclaceae genome was conducted using Orthofinder 2.5.4 (Emms and Kelly, 2019) with parameters -*evalue 1e-5, -seg yes, -soft_masking true, - use_sw_tback*. The results were filtered to the query coverage >= 75% and the percent identity >= 70%. Orthologous gene clusters were identified using MCL version 14-137 with an inflation value of 1.1 (Enright et al., 2002).

### 2.3 Phylogenetic analysis of pangenome

Orthofinder was used to identify the pan single-copy genes for reliable phylogenic tree construction and gene flux analysis. The pan single-copy genes were aligned using the linsi option in MAFFT version 7.508 (Katoh and Standley, 2013) and masked in Gblocks version 0.91b (Castresana, 2000). Seqkit (version 2.3.0, Shen et al., 2016) was used to sort the single copy gene sequences and convert the multi-line sequences into a one-line sequence. Iqtree version 2.2.0.3 (Minh et al., 2020) was used to predict the best phylogenetic tree model. Finally, the tree was constructed with model Q. insect+F+I+I+R4. Landscaping of the phylogenetic tree was achieved using iTOL version 6.6 (Letunic and Bork, 2021).

### 2.4 Pangenome analysis

When a genome set had incomplete genomes, it is necessary to determine a threshold number of genomes in which a gene must be observed in order to call it ‘core’. The probability that a gene was observed in all *Ca.* Accumulibacter genomes is the product of the completeness of each genome. The probability that a gene is missing in one genome but present in the rest genomes was calculated as the product of the completeness of the rest genomes multiplied by the incompleteness (i.e., 1 minus the completeness) of the incomplete genomes. Cut-off values were calculated using the R script (Zhang et al., 2019) (Supplementary Spreadsheet 1). The maximum number of genomes allowing an effective calculation of the cutoff value was 21. Via a comprehensive evaluation of the quality and the clade distribution of all available genomes, 21 high-quality *Ca.* Accumulibacter genomes covering 8 different clades were used for pangenomic analysis (The completeness and contamination of these genomes are documented in the Supplemental Materials Table S1-S2).

### 2.5 Gene gain/loss analysis

Gene flux was analyzed using Count (Csűös, 2010) based on the matrix of orthologous gene family abundance obtained in the foregoing analyses. Wagner parsimony penalty was set to 2 for better analysis of gene gain and loss (Pál et al., 2005; Zaremba-Niedzwiedzka et al., 2013). Genes acquired before the node of the last common ancestor (LCA) of *Ca.* Accumulibacter were defined as ancestral and genes acquired at the node of *Ca.* Accumulibacter LCA were defined as derived genes. Genes determined to be obtained via HGT in the derived genes were classified as laterally derived genes. Lineage specific genes were present in a single *Ca.* Accumulibacter genome. Flexible genes were present in more than one but less than 18 *Ca.* Accumulibacter genomes. Genetic comparisons were performed between PAO and GAO genomes to better understand the differences in their genetic makeup. The pangenome composed of 21 *Ca.* Accumulibacter and 2 *Dechloromonas* PAOs was denoted as the pan PAO genome. Core genes of the pan PAO genome were defined as genes belonged to the core genes of the pan *Ca.* Accumulibacter genome and were also present in two *Dechloromonas* PAO genomes. Differential genes were defined as core genes present in the pan PAO genome but absent in the *Ca.* Propionivibrio aalborgensis GAO genome.

### 2.6 Metabolic function analysis

The ancestral, derived, flexible and lineage-specific genes were annotated and classified based on KEGG annotations (Kanehisa et al., 2013) of clade IIC member SCUT-2 (Deng et al., 2023) and clade IIA member UW1 (García Martín et al., 2006; Oyserman et al., 2016b). The number of different types of genes annotated in each metabolic pathway was counted with the number of each type of genes being divided by the total number of genes in the pathway. Metabolic pathways with high proportions of derived genes were considered to have undergone major changes during evolution.

### 2.7 Horizontal gene transfer (HGT) identification

Parametric and phylogenetic methods are commonly used to infer HGT (Ravenhall et al., 2015). This study used the phylogenetic method for HGT identification. Each derived gene was queried in the NR database (published on May 7, 2015) (Pruitt et al., 2006) using the following BLASTP parameter [*-max_target_seqs 100-value 1E-6*] to preserve the first 100 BLAST results. The representative species were obtained from the first 100 BLAST results. The numbers and percentages of *Ca.* Accumulibacter, non-*Ca.* Accumulibacter Rhodocyclaceae, and non-Rhodocyclaceae members in the first 100 BLAST results were then calculated. A gene was considered as laterally derived gene if the numbers of *Ca.* Accumulibacter or non-*Ca.* Accumulibacter Rhodocyclaceae related hits were less than 10%. All core and differential derived genes in each metabolic pathway were analyzed to determine if they were obtained via HGT. The derived genes which were classified as HGT-originated are referred to as laterally derived genes. The origination of key genes (*ppk*2 and the homolog of *pho*U) was further confirmed using the phylogenetic method based on best-match analysis.

### 2.8 Metatranscriptomic analysis

An anaerobic-aerobic full cycle study was performed on an enrichment culture of *Ca.* Accumulibacter Clade IIC SCUT-2 in the lab-scale EBPR reactor SCUT (Supplementary Materials). The P cycling activities and the transformation of carbon compounds were monitored. Activated sludge samples were collected just before the start of a SBR cycle (0 min), and at 5 min (anaerobic phase), 30 min (anaerobic phase), 105 min (aerobic phase), and 120 min (aerobic phase) of the SBR cycle. The samples were snap-frozen in liquid N_2_, and stored at -80 °C before RNA extraction for metatranscriptomic analysis.

For metatranscriptomic analysis, total RNA was extracted using the RNA PowerSoil® Total RNA Isolation Kit (Omega Bio-Tek, GA, USA). Fastp (Chen et al., 2018) and SortMeRNA (Kopylova et al., 2012) were used to remove adaptation sequences and rRNAs. Filtered reads were mapped to the corresponding *Ca.* Accumulibacter draft genome (i.e., SCUT-2) using BBMap version 38.96 (Bushnell et al., 2017) and were normalized to transcript per million (TPM). Genes with TPM>100 were considered to be highly transcribed. Details on the reactor operation, full-cycle study, sample collection, metagenomic analysis and metatranscriptomic analysis are found in the Supplementary Materials. Raw reads and draft genomes obtained were submitted to NCBI under the BioProject No. PRJNA807832 and No. PRJNA771771.

## 3 Results

### 3.1 Identification of orthologous gene clusters

A total of 60,722 pan Rhodocyclaceae orthologous gene clusters were identified, including 25,080 homologous genes in the *Ca.* Accumulibacter pangenome (Supplementary Material Spreadsheet 2, Sheet 1 and 3). Large proportions (63.8% and 54.7%) of gene families in the pan Rhodocyclaceae and pan *Ca.* Accumulibacter genomes were present in only single genomes (Figure 2a and 2b). Approximately 1% (626) of gene families were present in >37 of the 43 genomes, which were used to define the core pan Rhodocyclaceae genome (Figure 2c). In the pan *Ca.* Accumulibacter genome, 6.9% genes were shared in >=18 genomes (Figure 2d). Non-paralogous genes (average gene copy per genome=1) account for high proportions of pan Rhodocyclaceae and pan *Ca.* Accumulibacter genomes (95.6% and 93.8%, respectively) (Figure 2e, 2f). The orthologous gene cluster identification results including the number of representative genes in each genome and summary statistics of pan Rhodocyclaceae and pan *Ca.* Accumulibacter gene clusters are provided in the Supplementary Material Spreadsheet 2 (Sheets 2 and 4).

**Figure 1.**
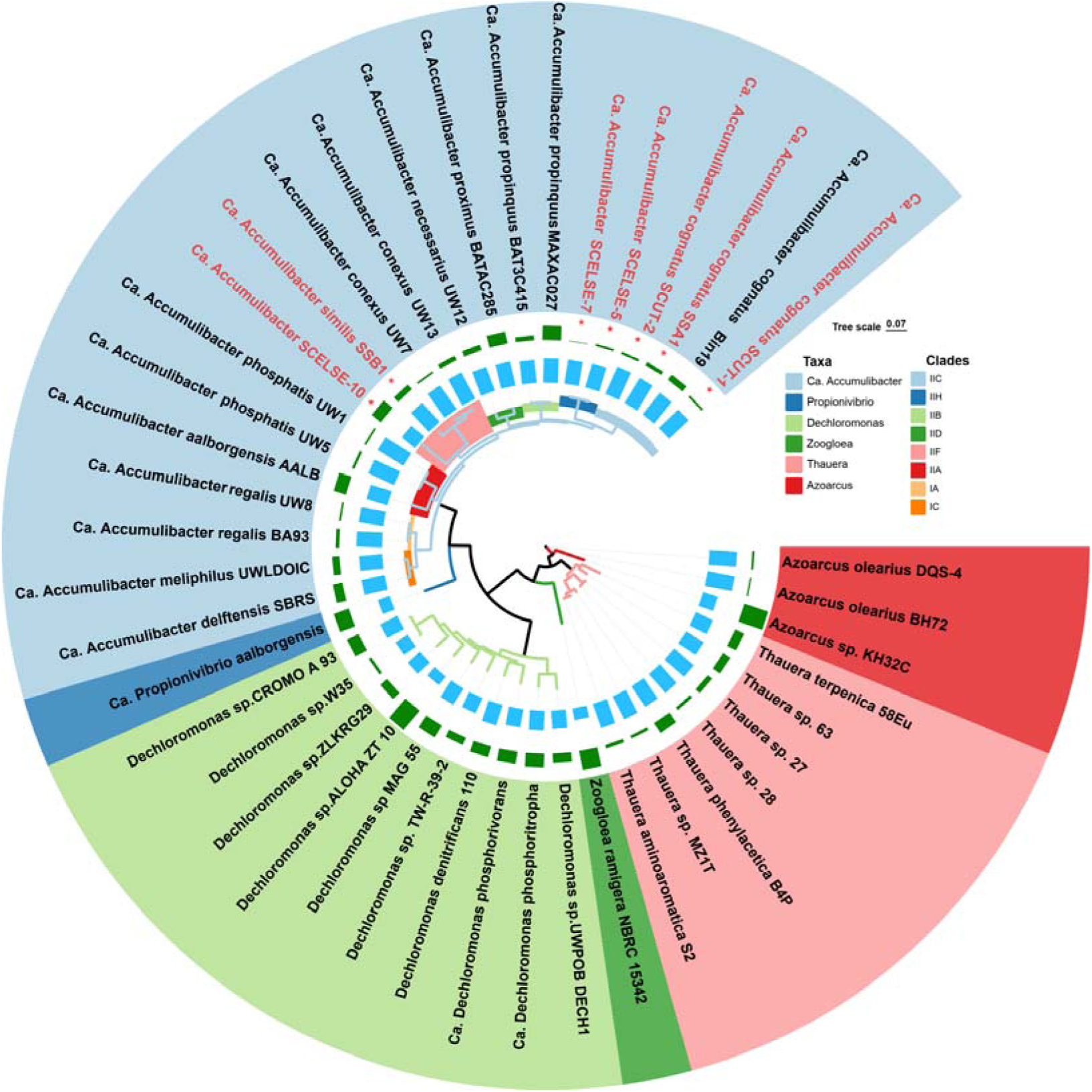
A phylogenetic tree of 43 Rhodocyclaceae members was built based on the concatenation of 59 single-copy genes. The genomes in red were recovered from our lab-scale reactors. SSA1, SSB1 and SCUT-1 were recovered in our previous work (Arumugam et al., 2019; Tian et al., 2022). SCELSE-5, SCELSE-7, SCELSE-10, and SCUT-2 were recovered from three of our lab-scale EBPR reactors (Supplementary Materials). The blue bars represent the number of shared orthogroups; and the green bars represent the number of unassigned genes.

**Figure 2.**
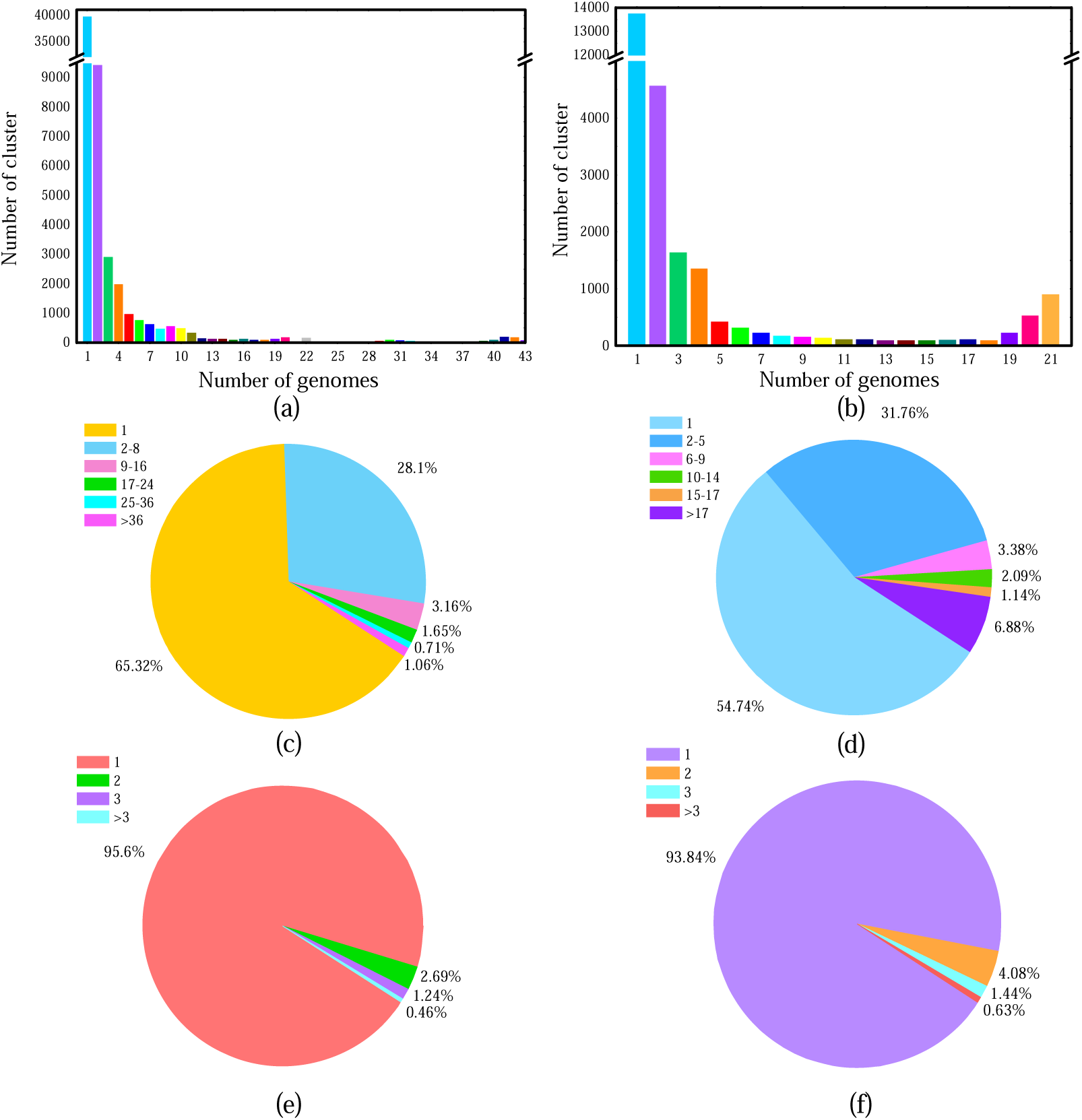
The number of gene clusters at different frequencies in (a) the pan Rhodocyclaceae genome and (b) the pan Accumulibacter genome. The proportion of clusters at different frequencies in (c) the pan Rhodocyclaceae genome and (d) the pan Accumulibacter genome. The proportion of different average gene copies per genome in (e) the pan Rhodocyclaceae genome and (f) the pan Accumulibacter genome. In each orthogroup, the average gene copies per genome are defined as the number of genes divided by the number of genomes.

### 3.2 Gene flux analysis

Among the 25,080 gene clusters in the pan *Ca.* Accumulibacter genome, 2499 (9.96%) were inferred to occur in the genome of the last common ancestor (LCA), and 1668 (6.73%) occurred before the LCA. Eight hundred eighteen (3.26%) were acquired at the node of LCA. Gene occurrence possibility calculation suggested that with a genome-number cutoff of 18, 99.94% of core genes could be identified (Figure 3a). At this cutoff value, 1725 (6.88%) core genes were identified in the pan *Ca.* Accumulibacter genome (Figure 3b and 3c). By further reducing the cutoff value to 17, the number of core genes increased from 1725 to 1829, and those with known functions increased from 298 to 318. As this study mainly focused on the changes in the genetic content, i.e., new core derived genes and horizontally transferred genes, looser cutoff values did not seem to bring new gains. Thus, a relatively stricter cut-off value (i.e., 18) was used to ensure the accuracy of the results. The gene gain or loss of a pangenome needs to be characterized in specific lineage member genomes. To facilitate a subsequent combination with the transcriptome data, SCUT-2 and UW1 were used as representative genomes for gene flux analysis. Each gene in Clade IIC SCUT-2 and Clade IIA UW1 genomes was classified as ancestral, derived, lineage-specific or flexible genes. There were no significant differences in the numbers and proportions of ancestral and flexible genes in these two genomes (ancestral genes accounted for 32.6% and 34.7%; flexible genes accounted for 43.8% and 43.6%, in SCUT-2 and UW-1, respectively). 638 and 802 derived genes were found in the SCUT-2 and UW1 genomes (17.6% and 14.0%, respectively). 189 lineage specific genes (genes occurred only in UW1) were observed in UW1, which was slightly less than those (i.e., 275) in the SCUT-2 genome (Figure 3d). Figure 4 and Supplementary Material Spreadsheet 3 provided additional details about the presence, gain and loss of genes, and the discrete categories to which they were assigned.

**Figure 3.**
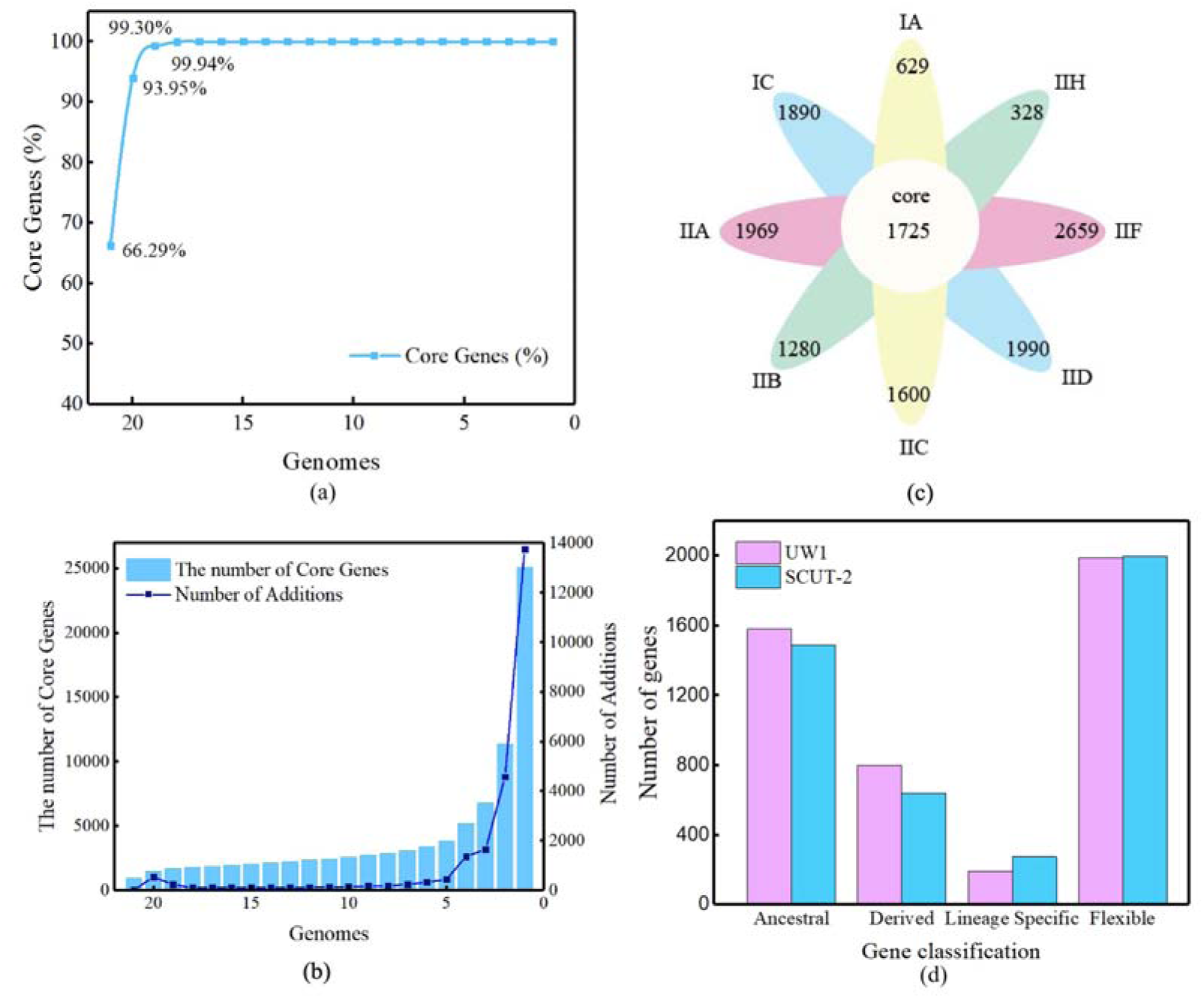
(a) Using the genome integrity estimate, about 99.94% of the core genes could be identified with a cut-off value of 18. Only gene families that appear in >=18 *Ca.* Accumulibacter genomes are considered as core genes. (b) The number of core genes observed at different cutoff values. (c) A Venn diagram describing the numbers of core genes and lineage-specific genes in the pan *Ca.* Accumulibacter genome. (d) The number of genes assigned as ancestral, derived, lineage-specific, and flexible genes in SCUT-2 and UW1.

**Figure 4.**
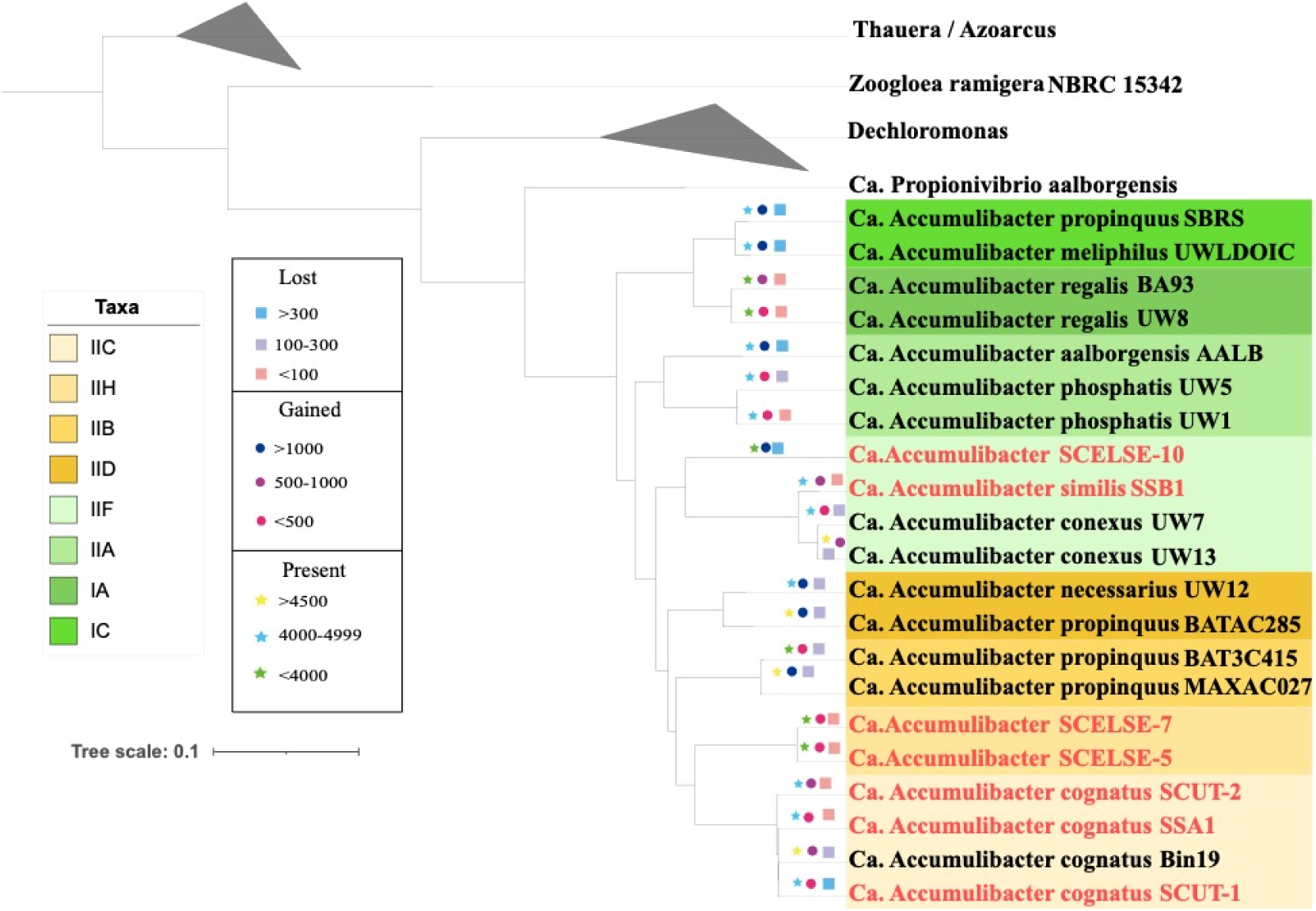
Gain or loss of genes at various nodes of the *Ca.* Accumulibacter lineage. A maximum likelihood tree was built based on the concatenation of single-copy genes with model Q. insect+F+I+I+R4. Genomes in red are those recovered from our bioreactors (Arumugam et al., 2019; Qiu et al., 2020; Tian et al., 2022; Deng et al., 2023).

### 3.3 Evolution of *Ca.* Accumulibacter metabolic pathways

The collections of genes identified as ancestral, derived, flexible, and lineage-specific genes were annotated using KEGG (Kanehisa et al., 2013) and were grouped into different metabolic pathways. In SCUT-2, 2293 genes were annotated to various metabolic pathways. The translation metabolic pathway had the highest proportion of ancestral genes (77, accounting for 96%). The largest number of ancestral genes (224) and derived genes (63) was observed in the carbohydrate metabolism pathway, accounting for 63% and 18%, respectively. The highest proportion (15 out of 53, 28.0%) of derived genes was observed in the cell growth and death metabolic pathway (Figure 5a). Within each primary pathway, ancestral and derived genes also showed distinct proportions in different secondary pathways. For instance, within the carbohydrate metabolism, the galactose metabolism pathways had the highest proportion (4 out of 5, 80%) of derived genes. Whereas, ancestral genes dominated the citric acid cycle (TCA cycle) (25 out of 30, 83%) and the glyoxylate and dicarboxylate metabolism pathways (33 out of 45, 73%). In signal transduction, the two-component system contained the highest proportion of derived genes (27 out of 182, 15%). In membrane transport, of the 122 ABC-transporter encoding genes, 18 were derived (15%) (Figure 5). Similar number and proportion of genes assigning to various metabolic pathway were observed in the *Ca*. Accumulibacter clade IIA UW1 genome with only two metabolic pathways (transport and catabolism, cell growth and death) showing significant difference in the proportions of derived genes (28%and 40% in SCUT-2 and 14% and 23% in UW1, respectively) (Figure 5 and Supplementary Material Figure S1). These results indicated that different strains of *Ca.* Accumulibacter underwent comparable developmental changes during evolution, but at the same time, preserved a certain degree of gene diversity. Detailed annotation of each gene in SCUT-2 and UW1 can be viewed in the Supplementary Spreadsheet 4.

**Figure 5.**
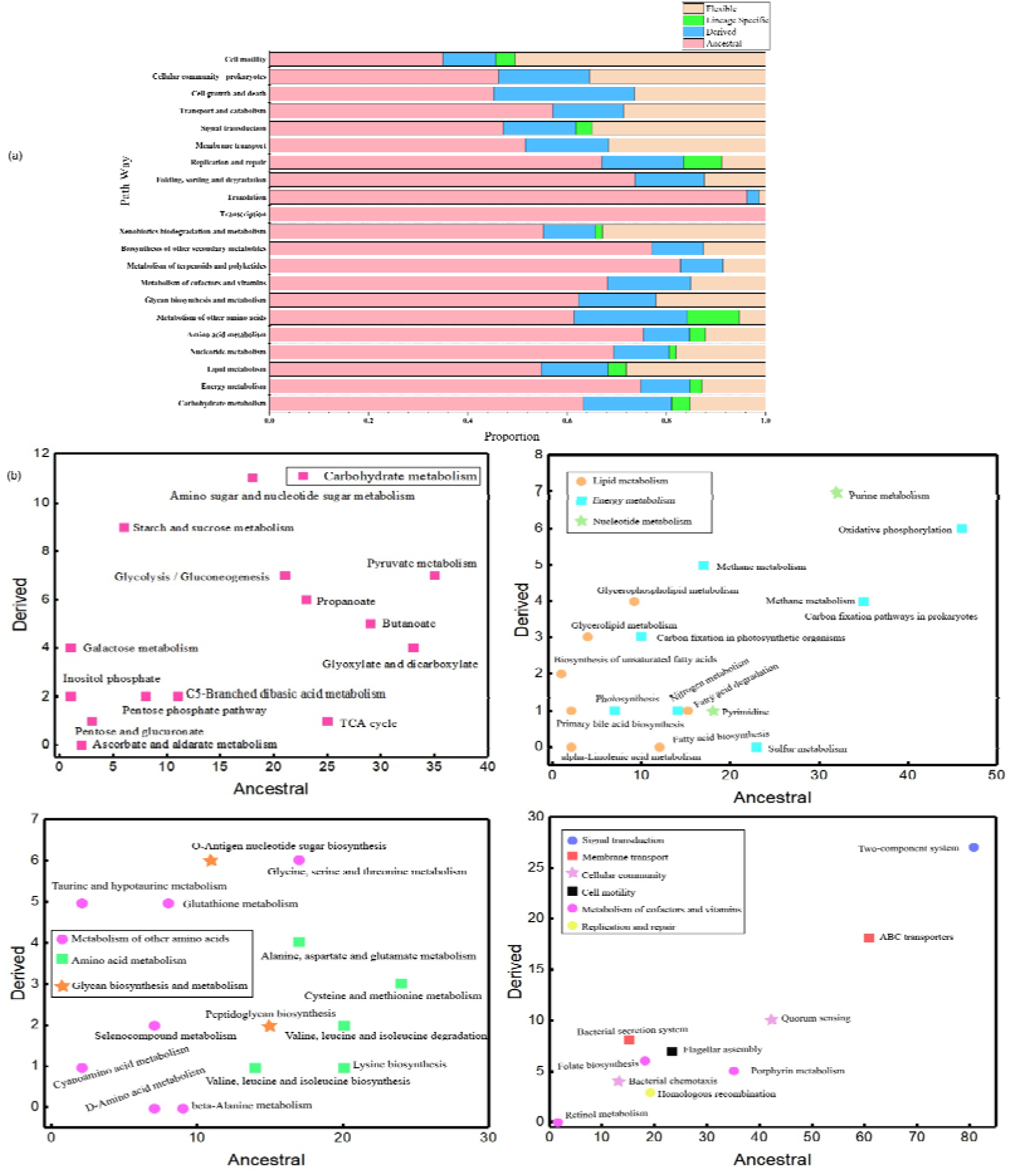
(a) The ratio of ancestral, flexible, lineage specific and derived genes in different primary metabolic pathways of SCUT-2. (b) Number of ancestral genes and derived genes in representative secondary pathways of SCUT-2.

### 3.4 Pan *Ca.* Accumulibacter phylogenetic analysis of derived genes

Relatively strict parameters (i.e., 70% identity and 75% coverage) were used to identify homologous gene clusters. The derived genes were manually classified into those derived from accumulative mutations and those from HGT. Phylogenetic analysis was further performed to confirm that *ppk*2 and the homolog of *pho*U are horizontally derived (Supplementary Materials Figure S4). Among 298 core derived genes that have been successfully annotated in KEGG, 124 were shown to have been acquired via HGT. Of the 124 genes, 67 were involved in KEGG pathways. The carbohydrate metabolism pathway harbors the highest numbers (25) of derived genes via HGT, including these in glycolysis/gluconeogenesis (e.g., genes encoding the phosphoglucomutase, the glucokinase, and the phosphoglycerate kinase), starch and sucrose metabolism (e.g., the starch synthase, and the glycogen phosphorylase genes), and in butanoate metabolism (genes encoding the poly[(R)-3-hydroxyalkanoate polymerase subunits). In signal transduction, the two-component system contained 10 laterally derived genes, such as genes encoding the REDOX signal transduction system proteins RegA/B, the phosphate regulon proteins PhoR-PhoB. Another remarkable set of genes derived via HGT were oxidative phosphorylation in the energy metabolism pathway including these encoding the NADH-quinone oxidoreductase subunit, the polyphosphate kinase, the cytochrome C. The inorganic phosphate transporter gene (*pit*) was also acquired via HGT. Similar results were observed for UW1. In the two-component system, genes encoding the REDOX signal transduction system proteins RegA/B, the phosphate regulon proteins PhoR-PhoB were laterally derived. More details about the BLAST comparison results can be found in the Supplementary Spreadsheet 5.

### 3.5 Comparison of genetic compositions in PAOs and non-PAOs

If a gene was present in *Ca.* Accumulibacter, but absent in other closely related PAOs, it may also not be a key to the development of the P cycling phenotype. On the other hand, if a gene was present in *Ca.* Accumulibacter or their closely related PAOs but absent in non-PAOs, it might be a key gene to the emergence of the PAO phenotype. For a better understanding of the genomic difference between closely related PAOs and non-PAOs, a pan PAO genome (Composed of 21 *Ca.* Accumulibacter and 2 *Dechloromonas* PAOs, Petriglieri et al., 2021) analysis was performed. The pan PAO genome was compared to the *Ca.* Propionivibrio aalborgensis (a closely related GAO, Albertsen et al., 2016) genome to identify differential genes (defined as core genes present in the pan PAO genome but absent in the *Ca.* Propionivibrio aalborgensis genome). In the pan PAO genome, 124 differential genes were identified. Alkaline phosphatase synthesis response regulator (PhoP) and polyphosphate kinase 2 (PPK2) genes were both differential genes. Other genes in the operon or the genes regulated by PhoP were not differential genes. Carbohydrate metabolism had the largest number of differential genes (16), including those encoding the acetyl-CoA C-acetyltransferase and the enoyl-CoA hydratase. The cofactor and vitamin metabolic pathway harbored the second largest number of differential genes (11), followed by energy metabolism (9), replication and repair (6) and signal transduction (5) metabolic pathways. The lowest number (1) of differential genes was observed in the transcription and metabolism of other amino acids pathways. A further analysis of another 21 available *Propionivibrio* genomes further confirmed that *ppk*2 and *pho*U are differential genes between *Ca.* Accumulibacter and *Propionivibrio*. HGT analysis was aimed at genes acquisition in *Ca.* Accumulibacter during evolution, based on the hypothesis that the emergence of the P cycling ability by PAOs was a result of acquisition of certain key genes. However, the hypothesis ignored the possibility that non-PAOs may have lost certain key genes in the process of evolution, leading to their inability to remove P. Differential genes included genes loss in non-PAOs during evolution. The analysis in this part allows us to more comprehensively understand the evolutionary process from a different perspective. More details about the differential genes (metabolic pathway and functional annotation) can be viewed in the Supplementary Spreadsheet 6.

### 3.6 Metatranscriptomic profiles

By analysis of the gene transcription levels of *Ca*. Accumulibacter in a typical EBPR cycle, genes which was not remarkably transcribed in the comparative genome may be excluded. Thus, the range of genes could be further narrowed down, facilitating the identification of key genes important to the PAO phenotype. Metatranscriptomic analysis was performed on an enrichment culture of *Ca*. Accumulibacter clade IIC strain SCUT-2 (with a relative abundance of 37.1% as suggested by the metagenomic analysis). In SCUT-2 genome, out of 5037 annotated genes, 906 were highly transcribed (TPM > 100). There were 298 core derived genes, 84 of which were highly transcribed (Supplementary Spreadsheet 7). To understand the dynamic patterns and functional relationships of 905 core genes with known function, they were classified into five clusters using the Mfuzz (Kumar, 2007) (Figure 6b). Most genes (e.g., the acetate permease gene *act*P, NOF05_02545) in Cluster 1 were related to transporter for carbon uptake and energy utilization. Cluster 2 showed a pattern of increased transcription throughout the anaerobic period, peaking after oxygen exposure. The phosphate transport system substrate binding protein (*pstS*, NOF05_04305) and the laterally derived polyphosphate kinase 2 gene (*ppk2*, NOF05_17285) showed Cluster 2 transcription pattern. Cluster 3 genes showed high transcription at the beginning of the anaerobic stage, and reduced towards the end of the anaerobic cycle as the depletion of acetate (Figure 6a). Their high transcription in the aerobic stage were mostly relate to the routing of anaerobically stored carbon to the TCA cycle and glycogenesis (Oyserman et al., 2016b; Qiu et al., 2020). Cluster 4 contained genes encoding the distant homolog of PhoU (NOF05_17860, NOF05_12350) and antitoxin CptB (NOF05_13125) which showed low transcription during the anaerobic stage but were upregulated during the aerobic phase, potentially playing a role in sustaining vital activities and controlling homeostatic environments (Shang et al., 2020). Genes in Cluster 5 may be associated with the maintenance of stable intracellular environments or cell growth, including genes encoding the ion transporters, such as the magnesium transporter gene (NOF05_18175) and the low affinity inorganic phosphate transporter (*pit*, NOF05_12345). These clustering patterns aligned with the metabolic characteristics of *Ca.* Accumulibacter in EBPR (Figure 6a).

**Figure 6.**
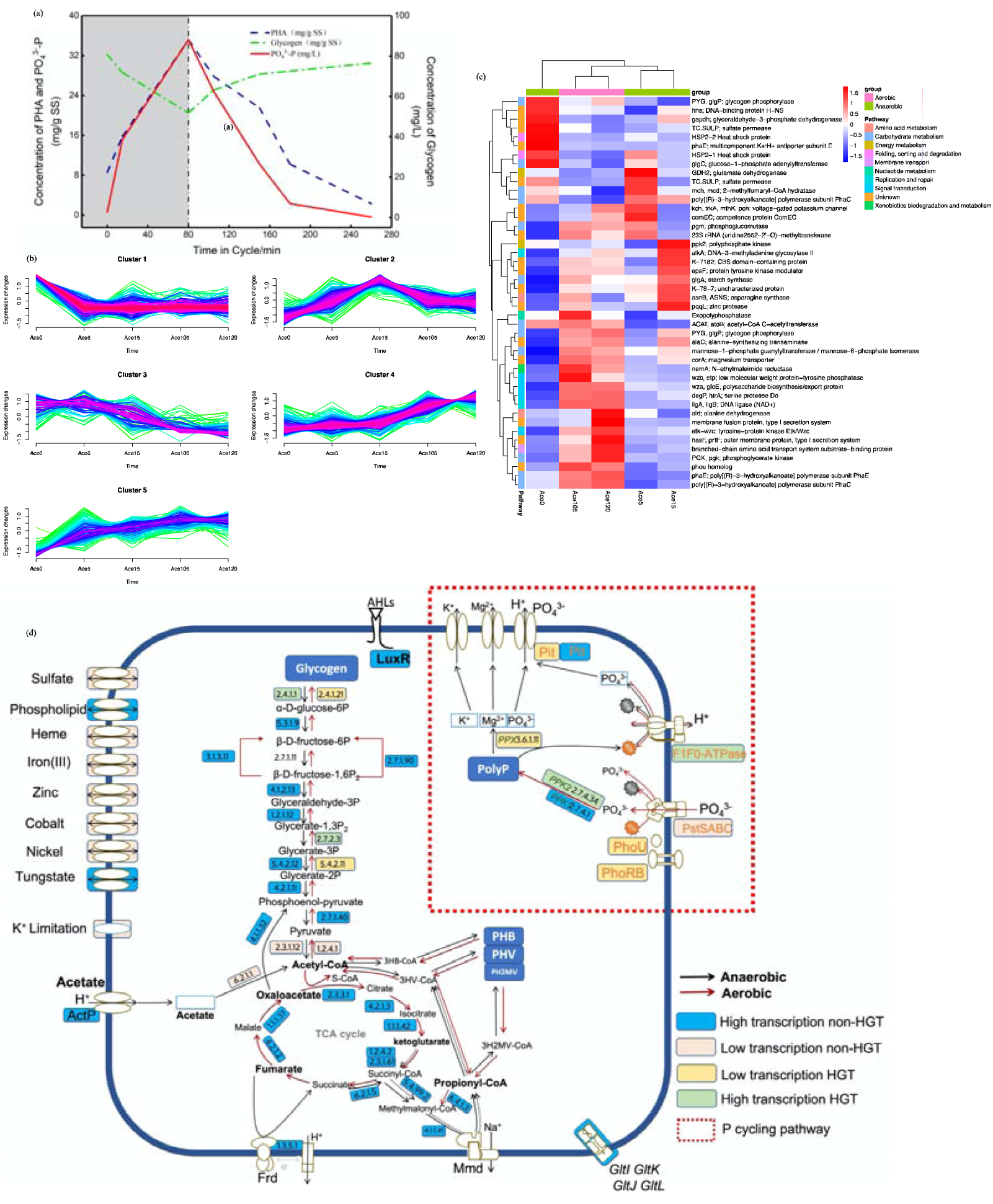
(a) Changes in phosphate, PHA and glycogen concentrations during an anaerobic-aerobic full cycle. (b) Cluster analysis of transcriptome data at different time points for transcription pattern identification. (c) 44 highly transcribed and laterally derived genes (via HGT) in the SCUT-2 genome during the anaerobic-aerobic full cycle. (d) A metabolic model of *Ca*. Accumulibacter. Black and red solid arrows represent active metabolic pathways in the anaerobic and aerobic phases, respectively. Genes in blue and pink are genes not acquired by HGT with high and low transcription, respectively. Genes in green and yellow represent genes acquired via HGT with high and low transcription, respectively. The red dashed line denotes the key P cycling pathway. The enzyme commission (EC) number indicates the key enzyme involved in each pathway/reaction.

The transcription of horizontally transferred genes in SCUT-2 was further analyzed. 44 genes, which were identified to be obtained via HGT, were highly transcribed (Figure 6c). These genes were involved in pathways, such as glycolysis/gluconeogenesis (phosphoglycerate kinase, and phosphoglucomutase), ABC transporters (branched-chain amino acid transport system substrate-binding protein), butanoate metabolism (poly[(R)-3-hydroxyalkanoate] polymerase subunit), two-component system (low molecular weight protein-tyrosine phosphatase, polysaccharide biosynthesis/export protein, tyrosine-protein kinase and serine protease), transporters for inorganic salts (sulfate permease, and magnesium transporter), and showed high transcription throughout the EBPR cycle. Polyphosphate kinase 2 gene (*ppk*2) was also highly transcribed and was significantly upregulated in the anaerobic phase. The transcription of phosphate transport regulator (a distant homolog of PhoU) was significantly upregulated in the aerobic stage. PHA synthesis related genes were also highly transcribed. A full list of the SCUT-2 gene transcription data can be found in the Supplementary Spreadsheet 7.

Comparisons were further made to the gene transcription characteristics of UW1 (Oyserman et al, 2016b). 35 horizontally derived gene families were high transcribed in both SCUT-2 and UW1 (Supplementary Material Figure S2). Apart from *pho*U and *pit* which are related to phosphate regulation and transport, 42 laterally derived gene families were under-transcribed in SCUT-2 but highly transcribed in UW1, including the acetate kinase gene. These 42 gene families may not play a key role in the evolution of non-PAO to PAO due to their different transcription behaviors in SCUT-2 and UW1. Combined with transcriptomic analysis, the range of key genes can be effectively reduced, and a metabolic model of *Ca*. Accumulibacter can be construct (Figure 6d). Most genes in the central carbon metabolic pathway were highly transcribed non-HGT genes, indicating that although this pathway is indispensable for *Ca.* Accumulibacter, its contribution to the evolution from a non-PAO metabolism to a PAO metabolism is unlikely. In the P cycling pathway, several laterally acquired genes were involved, potentially playing a key role in the evolution of *Ca*. Accumulibacter. Some of them were highly transcribed, potentially playing a key role in the evolution of *Ca.* Accumulibacter (Figure 6).

## 4 Discussion

Previous research suggested that the transition of PAO from non-PAO may have been occurred at the node of *Ca.* Accumulibacter LCA (Oyserman et al., 2016a). However, a recent study suggested that there are also PAOs in the *Dechloromonas* genus (i.e., *C*a. Dechloromonas phosphoritropha, *C*a. Dechloromonas phosphorivorans) (Petriglieri et al., 2021), raising a possibility that the emergence of the PAO phenotype may have occurred before the *Ca.* Accumulibacter LCA. Here we discuss the function of key laterally derived genes in the context of pangenomics and known PAO metabolism. A metatranscriptomic analysis of an enrichment culture of *Ca*. Accumulibacter Clade IIC member SCUT-2 contrasting those of *Ca*. Accumulibacter Clade IIA UW1 were performed to study the transcriptional dynamics of key genes in *Ca.* Accumulibacter. This approach allowed the exclusion of genes that were not highly transcribed in the large collection of laterally derived genes to narrow down the range of key genes to obtain new insights on key genomic features of the polyphosphate accumulating trait.

### 4.1 Carbon substrate uptake

The largest number of genes were annotated to the carbohydrate metabolism pathway in both SCUT-2 and UW1 genomes (354 and 369, respectively). In the SCUT-2 genome, there were 224 ancestral genes, 63 derived genes, and 49 laterally derived genes. Transcriptomic analysis suggested that, when acetate was used as a carbon source, genes directly related to intracellular acetate processing and PHA synthesis were remarkably upregulated in SCUT-2 (Supplementary Spreadsheet 7). The high affinity acetyl-CoA synthetase (NOF05_02565) and low-affinity phosphate acetyltransferase (NOF05_11790) are responsible for acetate activation (He et al., 2011). Other genes involved in the acetyl-CoA pathway, including the pyruvate kinase gene (NOF05_14290) and the phosphoenolpyruvate carboxykinase gene (NOF05_14615), maintained high levels of transcription throughout the anaerobic-aerobic cycle. However, these genes are all ancestral genes. There was only one horizontally transferred gene (i.e., the acetate kinase gene, NOF05_16845) which was barely transcribed. Therefore, genes related to acetate processing may not be keys to the emergence of the PAO phenotype. In addition, in the TCA cycle (Zhou et al., 2009), there were a total of 30 genes. Among them only the dihydrolipoamide dehydrogenase gene (NOF05_18520) was laterally derived, whereas transcribed at a low level, indicating that gain/loss of genes in the TCA cycle might have not contributed remarkably to the evolution of non-PAOs to PAOs. Four laterally derived genes occurred in the PHA synthesis pathway (*pha*C NOF05_18015, NOF05_21650, NOF05_21620 and *pha*A NOF05_18020), NOF05_21650 and NOF05_21620 were highly transcribed throughout the EBPR cycle (Figure 6). Whereas *Ca.* Propionivibrio aalborgensis also encoded these genes (Albertsen et al., 2016), their contribution to the evolution from a non-PAO metabolism to a PAO metabolism was unlikely.

### 4.2 Two-component systems

The two-component signal transduction system enables bacteria to sense, respond and adapt to diverse and dynamic environmental conditions (Capra and Laub, 2012). This system is commonly preserved in the bacterial domain. The number of genes in the two-component system was considered to be closely related to the bacteria’s living environment (Alm et al., 2006). Bacteria living in extreme environments tend to encode a large number of signaling proteins for improved adaption (Ulrich and Zhulin, 2010). In the SCUT-2 genome, a total of 182 genes were annotated to the two-component system, including 81 ancestral genes and 27 derived genes. 12 of them were laterally derived. In both SCUT-2 and UW1, phosphate regulon response regulator gene *pho*B (NOF05_18105), phosphate regulon sensor histidine kinase gene *pho*R (NOF05_18105) and redox signaling genes *reg*A and *reg*B (NOF05_11115, NOF05_11120) were laterally derived. RegB/RegA was shown to control and regulate a variety of basic metabolic processes in *Rhodobacter*, *Capsulatus* and *Sphaeroides*, such as photosynthesis, CO_2_ fixation, N_2_ assimilation, denitrification, and electron transport (Elsen et al., 2004) via direct or indirect control of respective operons (Dubbs et al., 2000; Elsen et al., 2000). However, both *reg*A and *reg*B were absent in two *Dechloromonas* PAO genomes (GCA_016722705.1, and GCA_016721185.1) (Petriglieri et al., 2021), suggesting that the redox signaling RegA/B were not indispensable for a PAO phenotype. PhoR-PhoB is present in both *Ca.* Accumulibacter and two *Dechloromonas* PAO genomes, which has the potential to play a role in PAO phenotype evolution. Since the PhoR-PhoB system is a part of the Pho regulon, further discussion was provided in the following subsection.

### 4.3 Phosphate regulatory system

The phosphate regulator (Pho) is a regulatory mechanism to maintain and manage inorganic phosphate concentrations in bacterial cells. The system typically consists of extracellular enzymes, transporters and enzymes involved in the intracellular storage of phosphate (Santos-Beneit, 2015). Signal transduction of Pho regulators requires seven proteins, including PhoR, PhoB, four components of the ABC transporter Pst (PstS, PstA, PstB, and PstC) and PhoU. An increase in the extracellular phosphate concentration near the PstSCAB transporter would result in increased binding of phosphate to PhoU, which would then inhibit the PhoR kinase activity and the PstSCAB transporter activity. In the absence of phosphate input, PhoU dissociates with phosphate, allowing the phosphate transport (Pst) to return to a normal working state (Choi et al., 2022). The above feedback control enables bacteria to maintain and control a relative stable intracellular phosphate concentration. Most of the genes in the Pho regulatory system in *Ca.* Accumulibacter are laterally derived, including those encoding PhoR and PhoB. In addition, three distant *pho*U homologs (NOF05_17860, NOF05_09930, NOF05_09935) were found in *Ca*. Accumulibacter genomes which are also horizontally acquired core genes. Distant homologs are pairs of proteins which have similar structures and functions but low gene sequence similarity (Monzon et al., 2022). The homolog *pho*U is located near *pit* in the *pit* operon in *Ca.* Accumulibacter genomes. Moreover, PhoR-PhoB is also present in two *Dechloromonas* PAO genomes (*Ca.* Dechloromonas phosphoritropha and *Ca.* Dechloromonas phosphorivorans).

In SCUT-2, the transcription of *pho*R (NOF05_18110) and *pho*B (NOF05_18105, NOF05_19100) was negligible. The transcription level of *pho*R (CAP2UW1_1997) in UW1 was also low. The transcription of *pho*B (CAP2UW1_1996) in the aerobic phase was slightly upregulated (with TPM values from 12 to 92) but was still at relatively low levels (Supplementary Spreadsheet 5) These results suggest that PhoR-PhoB in *Ca.* Accumulibacter were probably not active in perceiving phosphate concentrations. Similarly, the *pho*U genes were almost not transcribed (with the maximum TPM values <12, Figure 6). Although the homolog *pho*U genes showed high transcription, the trend was not in line with *pst*, indicating that the laterally derived PhoU or their homologs were not effectively regulating Pst (Supplementary Material Figure S3). The same phenomenon was observed in UW1 (Oehmen et al., 2007) and UW6 (McDaniel et al., 2021) metatranscriptome (Supplementary Spreadsheet 7). In *Staphylococcus aureus*, the absence of *pho*U homolog which located in the *pit* operon leads to the upregulation of phosphate transporter genes (*pst*), resulting in increased intracellular polyphosphate levels (Shang et al., 2020). In *Sinorhizobium meliloti*, the absence of *pho*U was shown to result in excessive accumulation of phosphate, which inactivate cells due to P poisoning, resulting in poor cell growth (diCenzo et al., 2017; Li and Zhang, 2007). Based on these results, two hypotheses are proposed. (1) PhoU in *Ca.* Accumulibacter was ineffective in regulating Pst even under high intracellular phosphate concentrations (no transcription of the *pho*U, and the unmatched transcription of *pho*U homolog and *pst*, Supplementary Material Figure S3). Pst continued to operate (as indicated by the high transcription of *pst* in the transcriptome, Supplementary Material Figure S3), resulting in excessive phosphate accumulation in cells (Figure 6a). The laterally derived PPK2 functioned (as suggested by the high transcription of *ppk*2, Supplementary Material Figure S3) to condense excess phosphate into poly-P to avoid P poisoning. The second is that, in *Ca*. Accumulibacter, since *pho*U, the homolog of *pho*U and *ppk*2 were laterally derived from different donor bacteria (*Rhodocyclace*, *Burkholderia* and *Gramaproteobacteria*, respectively, as suggested by the BLAST results, Supplementary SpreadSheet 5), their encoding proteins (i.e., PhoU, PhoU homologue and PPK2) may have incompatible phosphate activation/inactivation thresholds. PPK2 continued to synthesize poly-P by consuming intracellular phosphate transported via Pst, resulting in consistently low intracellular phosphate concentration, which was insufficient to combine with PhoU and/or its homologs to downregulate Pst. In the SCUT-2 and UW1 transcriptomes (Figure 6), PPK2 showed high levels of transcription during the entire EBPR cycle (with TPM values up to 12,481 in SCUT-2), which was further up-regulated in the aerobic stage, suggesting that PPK2 worked to synthesize poly-P by consuming phosphate which was imported via Pst, avoiding possible cell inactivation and poisoning due to elevated intracellular phosphate concentrations and achieved poly-P accumulation. In addition, *Ca.* Dechloromonas phosporitropha were lack of *pst*, *pho*U, *pho*B and *pho*R genes in the Pho regulon, which is consistent with our hypothesis that the Pho regulation may not work properly in PAOs. The transcriptomics data of *Microlunatus phosphovorus* (BioProject No. PRJNA984968) and proteomics data of *Tetrasphaera elongate* (obtained from Herbst et al., 2019) were further analyzed to check whether the same hypothesis could apply to other PAOs. In the transcriptome of *Microlunatus phosphovorus*, we found that the transcriptional patterns of *pst* were also inconsistent with those of *pho*U during an anaerobic and aerobic cycle (Supplementary Material). From the proteome of *Tetrasphaera elongate*, the relative abundances of Pst and PhoU did not vary significantly between anaerobic and oxic conditions; hence they were not significantly affected by changes in phosphate concentrations (Herbst et al., 2019). Taken together, these results suggest that in *Microlunatus phosphovorus* and *Tetrasphaera elongate*, the regulation of Pst by PhoU was not effective, and that the Pho dysregulation hypothesis may also apply to non-*Ca.* Accumulibacter PAOs. However, additional work is needed to confirm its broad applicability.

Despite that there is limited research on the Pho regulatory system in *Ca.* Accumulibacter, the transcriptomics and gene origination analysis in the Pho regulon suggested that it may represent a key link in the emergence of the PAO phenotype.

### 4.4 Transport of phosphate

Phosphorus (organic and/or inorganic) are typical restricting nutrients. Therefore, microorganisms developed adaptive mechanisms to cope with ordinary P deficiency. Low affinity inorganic phosphate transport system (Pit) and high affinity phosphate transport system (Pst) are key transporters used for inorganic phosphate transport (Willsky and Malamy, 1980; Martín and Liras, 2021). In the pan *Ca.* Accumulibacter genomes, genes encoding the Pst transporter are not core genes, nor laterally derived genes. Furthermore, *Ca.* Dechloromonas phosporitropha (PAO) do not seem to encode any *pst* (Petriglieri et al., 2021). These results suggested that the Pst transport system may not be indispensable for a PAO phenotype. *Ca.* Dechloromonas phosporitropha encoded a phosphonates/phosphate transport system (Phn), which was shown to be a high affinity phosphate transporter in *Mycobacterium smegmatis* (Gebhard et al., 2006). This system may serve as a backup for the Pst transport system in *Ca.* Dechloromonas phosporitropha.

In the pan PAO genome, the low affinity inorganic phosphate transporter gene (*pit*, NOF05_09925, NOF05_09940) was laterally derived gene. The efflux of phosphate in symport with H^+^ via Pit results in the production of proton motive force, which is a key drive force for the uptake of VFAs and amino acid by *Ca.* Accumulibacter (Saunders et al., 2007; Qiu et al., 2020, Chen et al., 2022). Therefore, *pit* is an important feature gene for the PAO phenotype. In SCUT-2 transcriptomes, the transcription of the *pit* was upregulated during the transition from anaerobic to aerobic conditions (Supplementary Material Figure S3). The confirmed GAO, *Ca.* Propionivibrio aalborgensis, which are closely related to *Ca.* Accumulibacter (Figure 1), are lack of *pit*. But *pit* is present in the genomes of other GAOs, for example, *Defluviicoccus* GAO-HK (Wang et al., 2014), *Ca.* Competibacter denitrificans, and *Ca.* Contendobacter odensis (McIlroy et al., 2014). In addition, we analyzed 21 *Propionivibrio* genomes in the NCBI database. Pit transporter was encoded in 13 of 21 *Propionivibrio* genomes (Supplementary Material Table S4). Anyhow, *pit* may not be a key feature driven the evolution of non-PAO into PAOs and may neither be used as a marker gene for the PAO phenotype, although it is indispensable for the P cycling trait.

## 5. Conclusion

Pangomics in combination of a metatranscriptomic analysis of an enrichment culture of *Ca.* Accumulibacter clade IIC member SCUT-2 was performed to understand the genomic transition in the evolution of *Ca.* Accumulibacter, and to identify the key genes to the emergence of the P accumulating traits.

(1) 298 core genes were shown to have newly obtained by *Ca.* Accumulibacter at their least common ancestor. 124 of them were derived via HGT. 44 of these laterally derived core genes were highly transcribed in a typical EBPR cycle.

(2) High affinity phosphate transport system (Pst) may not be indispensable for the PAO phenotype. Inorganic phosphate transporter (Pit) may not be a key feature driving the evolution of non-PAO into PAOs. Their encoding genes may neither be used as a marker gene for the PAO phenotype.

(3) Low transcription of the *pho*R-*pho*B two-component system genes and the unmatched transcription of *pst* and *pho*U implied that the Pho regulon was likely not well functioning in *Ca.* Accumulibacter.

(4) A Pho dysregulation hypothesis was proposed. The laterally derived PhoU in *Ca.* Accumulibacter was not effective in regulating the Pst, resulting in excessive P uptake. To avoid P poisoning, the laterally derived PPK2 was employed to condense excess phosphate into poly-P. Alternatively, PhoU and PPK2 genes were derived from different donor bacteria, resulting in their unmatched activation/inactivation thresholds. PPK2 tend to reduce the intracellular phosphate to concentration levels which was perceived by PhoU as low-phosphate states, resulting in continuous phosphate uptake.

This study is expected to provide a new perspective for the understanding of the development and evolution of the P cycling traits for *Ca.* Accumulibacter.

## Declaration of Competing Interest

The authors declare that they have no known competing financial interests or personal relationships that could have appeared to influence the work reported in this paper.

## Acknowledgments

This research was supported by the National Natural Science Foundation of China (52270035 and 51808297), the Natural Science Foundation of Guangdong Province (2021A1515010494), the Guangzhou Key Research and Development Program (2023B03J1334), and the Pearl River Talent Recruitment Program (2019QN01L125).

## Data available

All data generated or analyzed during this study are included in this published article. Metagenomic raw reads and draft genomes were submitted to NCBI under the BioProject No. PRJNA807832 and No. PRJNA771771. Metatranscriptomic data were submitted to NCBI under the submitted No. PRJNA807832. Other data were documented in the Supplementary Materials.

## CRediT authorship contribution statement

**Xiaojing Xie:** Conceptualization, Methodology, Software, Formal analysis, Investigation, Data Curation, Writing – Original Draft, Visualization.

**Xuhan Deng:** Data Curation, Resources, Visualization.

**Liping Chen:** Data Curation, Resources, Visualization.

**Jing Yuan:** Investigation, Resources, Data Curation.

**Hang Chen:** Investigation, Resources, Data Curation.

**Chaohai Wei:** Writing – review & editing, Supervision.

**Xianghui Liu:** Investigation, Resources, Data Curation.

**Stefan Wuertz:** Supervision, Writing – review & editing, Project administration, Funding acquisition.

**Guanglei Qiu:** Conceptualization, Methodology, Investigation, Supervision, Writing – review & editing, Validation, Project administration, Funding acquisition.

## Notes

### Competing Interest Statement

The authors have declared no competing interest.

## Reference

Abdelfattah, A., Ali, S.S., Ramadan, H., El-Aswar, E.I., Eltawab, R., Ho, S.-H., Elsamahy, T., Li, S., El-Sheekh, M.M., Schagerl, M., Kornaros, M., Sun, J., 2023. Microalgae-based wastewater treatment: Mechanisms, challenges, recent advances, and future prospects. Environmental Science and Ecotechnology 13, 100205.

Aggarwal, S.K., Singh, A., Choudhary, M., Kumar, A., Rakshit, S., Kumar, P., Bohra, A., Varshney, R.K., 2022. Pangenomics in microbial and crop research: Progress, applications, and perspectives. Genes 13 (4), 598.

Albertsen, M., McIlroy, S.J., Stokholm-Bjerregaard, M., Karst, S.M., Nielsen, P.H., 2016. “*Candidatus* Propionivibrio aalborgensis”: A novel glycogen accumulating organism abundant in full-scale enhanced biological phosphorus removal plants. Frontiers in Microbiology 7.

Alm, E., Huang, K., Arkin, A., 2006. The Evolution of two-component systems in bacteria reveals different strategies for niche adaptation. PLoS Computational Biology 2 (11), e143.

Arumugam, K., Bağcı, C., Bessarab, I., Beier, S., Buchfink, B., Górska, A., Qiu, G., Huson, D.H., Williams, R.B.H., 2019. Annotated bacterial chromosomes from frame-shift-corrected long-read metagenomic data. Microbiome 7 (1), 61.

Bessarab, I., Maszenan, A.M., Haryono, M.A.S., Arumugam, K., Saw, N., Seviour, R.J., Williams, R.B.H., 2022. Comparative genomics of members of the genus *Defluviicoccus* with insights into their ecophysiological importance. Front Microbiol 13, 834906.

Bunce, J.T., Ndam, E., Ofiteru, I.D., Moore, A., Graham, D.W., 2018. A review of phosphorus removal technologies and their applicability to small-scale domestic wastewater treatment systems. Frontiers in Environmental Science 6, 8.

Bushnell, B., Rood, J., Singer, E., 2017. BBMerge – Accurate paired shotgun read merging via overlap. PLoS One 12 (10), e0185056.

Camejo Pamela, Y., Oyserman Ben, O., McMahon Katherine, D., Noguera Daniel, R., 2019. Integrated omic analyses provide evidence that a “*Candidatus* Accumulibacter phosphatis” strain performs denitrification under microaerobic conditions. mSystems 4 (1), e00193–00118.

Capra, E.J., Laub, M.T., 2012. Evolution of two-component signal transduction systems. Annual Review of Microbiology 66, 325–347.

Castresana, J., 2000. Selection of conserved blocks from multiple alignments for their use in phylogenetic analysis. Molecular Biology and Evolution 17 (4), 540–552.

Chen, L., Chen, H., Hu, Z., Tian, Y., Wang, C., Xie, P., Deng, X., Zhang, Y., Tang, X., Lin, X., Li, B., Wei, C.,Qiu, G. 2022. Carbon uptake bioenergetics of PAOs and GAOs in full-scale enhanced biological phosphorus removal systems. Water Research 216, 118258.

Chen, S., Zhou, Y., Chen, Y., Gu, J. 2018. fastp: an ultra-fast all-in-one FASTQ preprocessor. Bioinformatics 34 (17), i884–i890.

Choi, S., Jeong, G., Choi, E., Lee, E.-J., 2022. A dual regulatory role of the PhoU protein in *Salmonella Typhimurium*. mBio 13 (3), e00811–00822.

Coghlan, A., Tyagi, R., Cotton, J.A., Holroyd, N., Rosa, B.A., Tsai, I.J., Laetsch, D.R., Beech, R.N., Day, T.A., Hallsworth-Pepin, K., Ke, H.-M., Kuo, T.-H., Lee, T.J., Martin, J., Maizels, R.M., Mutowo, P., Ozersky, P., Parkinson, J., Reid, A.J., Rawlings, N.D., Ribeiro, D.M., Swapna, L.S., Stanley, E., Taylor, D.W., Wheeler, N.J., Zamanian, M., Zhang, X., Allan, F., Allen, J.E., Asano, K., Babayan, S.A., Bah, G., Beasley, H., Bennett, H.M., Bisset, S.A., Castillo, E., Cook, J., Cooper, P.J., Cruz-Bustos, T., Cuéllar, C., Devaney, E., Doyle, S.R., Eberhard, M.L., Emery, A., Eom, K.S., Gilleard, J.S., Gordon, D., Harcus, Y., Harsha, B., Hawdon, J.M., Hill, D.E., Hodgkinson, J., Horák, P., Howe, K.L., Huckvale, T., Kalbe, M., Kaur, G., Kikuchi, T., Koutsovoulos, G., Kumar, S., Leach, A.R., Lomax, J., Makepeace, B., Matthews, J.B., Muro, A., O’Boyle, N.M., Olson, P.D., Osuna, A., Partono, F., Pfarr, K., Rinaldi, G., Foronda, P., Rollinson, D., Samblas, M.G., Sato, H., Schnyder, M., Scholz, T., Shafie, M., Tanya, V.N., Toledo, R., Tracey, A., Urban, J.F., Wang, L.-C., Zarlenga, D., Blaxter, M.L., Mitreva, M., Berriman, M., International Helminth Genomes, C., 2019. Comparative genomics of the major parasitic worms. Nature Genetics 51 (1), 163–174.

Csűös, M., 2010. Count: evolutionary analysis of phylogenetic profiles with parsimony and likelihood. Bioinformatics 26 (15), 1910–1912.

Della Coletta, R., Qiu, Y., Ou, S., Hufford, M.B., Hirsch, C.N., 2021. How the pan-genome is changing crop genomics and improvement. Genome Biology 22 (1), 3.

Deng, X., Yuan, J., Chen, L., Chen, H., Wei, C., Nielsen, P.H., Wuertz, S., Qiu, G., 2023. CRISPR-Cas phage defense systems and prophages in *Candidatus* Accumulibacter. Water Research 235, 119906.

Diaz, R., Mackey, B., Chadalavada, S., kainthola, J., Heck, P., Goel, R., 2022. Enhanced Bio-P removal: Past, present, and future – A comprehensive review. Chemosphere 309, 136518.

diCenzo, G.C., Sharthiya, H., Nanda, A., Zamani, M., Finan, T.M., 2017. PhoU allows rapid adaptation to high phosphate concentrations by modulating PstSCAB transport rate in *Sinorhizobium meliloti*. Journal of Bacteriology 199 (18).

Dorofeev, A.G., Nikolaev, Y.A., Mardanov, A.V., Pimenov, N.V., 2020. Role of phosphate-accumulating bacteria in biological phosphorus removal from wastewater. Applied Biochemistry and Microbiology 56 (1), 1–14.

Dubbs, J.M., Bird, T.H., Bauer, C.E., Tabita, F.R., 2000. Interaction of CbbR and RegA* transcription regulators with the *Rhodobacter sphaeroides* cbbI Promoter-operator region. Journal of Biological Chemistry 275 (25), 19224–19230.

El-Sayed, N.M., Myler, P.J., Blandin, G., Berriman, M., Crabtree, J., Aggarwal, G., Caler, E., Renauld, H., Worthey, E.A., Hertz-Fowler, C., Ghedin, E., Peacock, C., Bartholomeu, D.C., Haas, B.J., Tran, A.-N., Wortman, J.R., Alsmark, U.C.M., Angiuoli, S., Anupama, A., Badger, J., Bringaud, F., Cadag, E., Carlton, J.M., Cerqueira, G.C., Creasy, T., Delcher, A.L., Djikeng, A., Embley, T.M., Hauser, C., Ivens, A.C., Kummerfeld, S.K., Pereira-Leal, J.B., Nilsson, D., Peterson, J., Salzberg, S.L., Shallom, J., Silva, J.C., Sundaram, J., Westenberger, S., White, O., Melville, S.E., Donelson, J.E., Andersson, B., Stuart, K.D., Hall, N., 2005. Comparative genomics of trypanosomatid parasitic protozoa. Science 309 (5733), 404–409.

Elsen, S., Dischert, W., Colbeau, A., Bauer, C.E., 2000. Expression of uptake hydrogenase and molybdenum nitrogenase in *Rhodobacter capsulatus* is coregulated by the RegB-RegA two-component regulatory system. Journal of Bacteriology 182 (10), 2831–2837.

Elsen, S., Swem, L.R., Swem, D.L., Bauer, C.E., 2004. RegB/RegA, a highly conserved redox-responding global two-component regulatory system. Microbiology and Molecular Biology Review 68 (2), 263–279.

Emms, D.M., Kelly, S., 2019. OrthoFinder: phylogenetic orthology inference for comparative genomics. Genome Biology 20 (1), 238.

Enright, A.J., Van Dongen, S., Ouzounis, C.A., 2002. An efficient algorithm for large-scale detection of protein families. Nucleic Acids Res 30 (7), 1575–1584.

Feng, S., Stiller, J., Deng, Y., Armstrong, J., Fang, Q., Reeve, A.H., Xie, D., Chen, G., Guo, C., Faircloth, B.C., Petersen, B., Wang, Z., Zhou, Q., Diekhans, M., Chen, W., Andreu-Sánchez, S., Margaryan, A., Howard, J.T., Parent, C., Pacheco, G., Sinding, M.-H.S., Puetz, L., Cavill, E., Ribeiro, Â.M., Eckhart, L., Fjeldså, J., Hosner, P.A., Brumfield, R.T., Christidis, L., Bertelsen, M.F., Sicheritz-Ponten, T., Tietze, D.T., Robertson, B.C., Song, G., Borgia, G., Claramunt, S., Lovette, I.J., Cowen, S.J., Njoroge, P., Dumbacher, J.P., Ryder, O.A., Fuchs, J., Bunce, M., Burt, D.W., Cracraft, J., Meng, G., Hackett, S.J., Ryan, P.G., Jønsson, K.A., Jamieson, I.G., da Fonseca, R.R., Braun, E.L., Houde, P., Mirarab, S., Suh, A., Hansson, B., Ponnikas, S., Sigeman, H., Stervander, M., Frandsen, P.B., van der Zwan, H., van der Sluis, R., Visser, C., Balakrishnan, C.N., Clark, A.G., Fitzpatrick, J.W., Bowman, R., Chen, N., Cloutier, A., Sackton, T.B., Edwards, S.V., Foote, D.J., Shakya, S.B., Sheldon, F.H., Vignal, A., Soares, A.E.R., Shapiro, B., González-Solís, J., Ferrer-Obiol, J., Rozas, J., Riutort, M., Tigano, A., Friesen, V., Dalén, L., Urrutia, A.O., Székely, T., Liu, Y., Campana, M.G., Corvelo, A., Fleischer, R.C., Rutherford, K.M., Gemmell, N.J., Dussex, N., Mouritsen, H., Thiele, N., Delmore, K., Liedvogel, M., Franke, A., Hoeppner, M.P., Krone, O., Fudickar, A.M., Milá, B., Ketterson, E.D., Fidler, A.E., Friis, G., Parody-Merino, Á.M., Battley, P.F., Cox, M.P., Lima, N.C.B., Prosdocimi, F., Parchman, T.L., Schlinger, B.A., Loiselle, B.A., Blake, J.G., Lim, H.C., Day, L.B., Fuxjager, M.J., Baldwin, M.W., Braun, M.J., Wirthlin, M., Dikow, R.B., Ryder, T.B., Camenisch, G., Keller, L.F., DaCosta, J.M., Hauber, M.E., Louder, M.I.M., Witt, C.C., McGuire, J.A., Mudge, J., Megna, L.C., Carling, M.D., Wang, B., Taylor, S.A., Del-Rio, G., Aleixo, A., Vasconcelos, A.T.R., Mello, C.V., Weir, J.T., Haussler, D., Li, Q., Yang, H., Wang, J., Lei, F., Rahbek, C., Gilbert, M.T.P., Graves, G.R., Jarvis, E.D., Paten, B., Zhang, G., 2020. Dense sampling of bird diversity increases power of comparative genomics. Nature 587 (7833), 252–257.

Fernandez-Fueyo, E., Ruiz-Dueñas, F.J., Ferreira, P., Floudas, D., Hibbett, D.S., Canessa, P., Larrondo, L.F., James, T.Y., Seelenfreund, D., Lobos, S., Polanco, R., Tello, M., Honda, Y., Watanabe, T., Watanabe, T., Ryu, J.S., Kubicek, C.P., Schmoll, M., Gaskell, J., Hammel, K.E., St. John, F.J., Vanden Wymelenberg, A., Sabat, G., Splinter BonDurant, S., Syed, K., Yadav, J.S., Doddapaneni, H., Subramanian, V., Lavín, J.L., Oguiza, J.A., Perez, G., Pisabarro, A.G., Ramirez, L., Santoyo, F., Master, E., Coutinho, P.M., Henrissat, B., Lombard, V., Magnuson, J.K., Kües, U., Hori, C., Igarashi, K., Samejima, M., Held, B.W., Barry, K.W., LaButti, K.M., Lapidus, A., Lindquist, E.A., Lucas, S.M., Riley, R., Salamov, A.A., Hoffmeister, D., Schwenk, D., Hadar, Y., Yarden, O., de Vries, R.P., Wiebenga, A., Stenlid, J., Eastwood, D., Grigoriev, I.V., Berka, R.M., Blanchette, R.A., Kersten, P., Martinez, A.T., Vicuna, R., Cullen, D., 2012. Comparative genomics of *Ceriporiopsis subvermispora* and *Phanerochaete chrysosporium* provide insight into selective ligninolysis. Proceedings of the National Academy of Sciences 109 (14), 5458–5463.

Fernández-Gómez, B., Richter, M., Schüler, M., Pinhassi, J., Acinas, S.G., González, J.M., Pedrós-Alió, C., 2013. Ecology of marine Bacteroidetes: a comparative genomics approach. The ISME Journal 7 (5), 1026–1037.

Flowers, J.J., He, S., Malfatti, S., del Rio, T.G., Tringe, S.G., Hugenholtz, P., McMahon, K.D., 2013. Comparative genomics of two ’*Candidatus* Accumulibacter’ clades performing biological phosphorus removal. The ISME Journal 7 (12), 2301–2314.

Gebhard, S., Tran, S.L., Cook, G.M., 2006. The Phn system of *Mycobacterium smegmatis*: a second high-affinity ABC-transporter for phosphate. Microbiology 152 (11), 3453–3465.

García Martín, H.G., Ivanova, N., Kunin, V., Warnecke, F., Barry, K.W., McHardy, A.C., Yeates, C., He, S., Salamov, A.A., Szeto, E., Dalin, E., Putnam, N.H., Shapiro, H.J., Pangilinan, J.L., Rigoutsos, I., Kyrpides, N.C., Blackall, L.L., McMahon, K.D.,Hugenholtz, P. 2006. Metagenomic analysis of two enhanced biological phosphorus removal (EBPR) sludge communities. Nature Biotechnology 24 (10), 1263–1269.

Golicz, A.A., Batley, J., Edwards, D., 2016. Towards plant pangenomics. Plant Biotechnology Journal 14 (4), 1099–1105.

He, S., McMahon, K.D., 2011. Microbiology of ’*Candidatus* Accumulibacter’ in activated sludge. Microbial Biotechnology 4 (5), 603–619.

Herbst, F.A., Dueholm, M.S., Wimmer, R., Nielsen, P.H., 2019. The proteome of *Tetrasphaera elongata* is adapted to changing conditions in wastewater treatment plants. Proteomes 7 (2).

Kanehisa, M., Goto, S., Sato, Y., Kawashima, M., Furumichi, M., Tanabe, M., 2013. Data, information, knowledge and principle: back to metabolism in KEGG. Nucleic Acids Research 42 (D1), D199–D205.

Katoh, K., Standley, D.M., 2013. MAFFT Multiple sequence alignment software version 7: Improvements in performance and usability. Molecular Biology and Evolution 30 (4), 772–780.

Kjærbølling, I., Vesth, T., Frisvad, J.C., Nybo, J.L., Theobald, S., Kildgaard, S., Petersen, T.I., Kuo, A., Sato, A., Lyhne, E.K., Kogle, M.E., Wiebenga, A., Kun, R.S., Lubbers, R.J.M., Mäkelä, M.R., Barry, K., Chovatia, M., Clum, A., Daum, C., Haridas, S., He, G., LaButti, K., Lipzen, A., Mondo, S., Pangilinan, J., Riley, R., Salamov, A., Simmons, B.A., Magnuson, J.K., Henrissat, B., Mortensen, U.H., Larsen, T.O., de Vries, R.P., Grigoriev, I.V., Machida, M., Baker, S.E., Andersen, M.R., 2020. A comparative genomics study of 23 *Aspergillus* species from section Flavi. Nature Communications 11 (1), 1106.

Kolakovic, S., Freitas, E.B., Reis, M.A.M., Carvalho, G., Oehmen, A., 2021. Accumulibacter diversity at the sub-clade level impacts enhanced biological phosphorus removal performance. Water Research 199, 117210.

Kopylova, E., Noé, L., Touzet, H., 2012. SortMeRNA: fast and accurate filtering of ribosomal RNAs in metatranscriptomic data. Bioinformatics 28 (24), 3211–3217.

Kumar, L., M, E.F., 2007. Mfuzz: a software package for soft clustering of microarray data. Bioinformation 2 (1), 5–7.

Letunic, I., Bork, P., 2021. Interactive Tree Of Life (iTOL) v5: an online tool for phylogenetic tree display and annotation. Nucleic Acids Research 49 (W1), W293–W296.

Li, Y., Zhang, Y., 2007. PhoU is a persistence switch involved in persister formation and tolerance to multiple antibiotics and stresses in *Escherichia coli*. Antimicrob Agents Chemother 51 (6), 2092–2099.

Liu, M., Huang, Y., Hu, J., He, J., Xiao, X., 2023. Algal community structure prediction by machine learning. Environmental Science and Ecotechnology 14, 100233.

Mao, Y., Graham, D.W., Tamaki, H., Zhang, T., 2015. Dominant and novel clades of *Candidatus* Accumulibacter phosphatis in 18 globally distributed full-scale wastewater treatment plants. Scientific Reports 5 (1), 11857.

Maszenan, A.M., Bessarab, I., Williams, R.B.H., Petrovski, S., Seviour, R.J., 2022. The phylogeny, ecology and ecophysiology of the glycogen accumulating organism (GAO) *Defluviicoccus* in wastewater treatment plants. Water Research 221, 118729.

Martín, J.F., Liras, P., 2021. Molecular mechanisms of phosphate sensing, transport and signalling in *Streptomyces* and related actinobacteria. International Journal of Molecular Sciences 22 (3), 1129.

McDaniel, E.A., Moya-Flores, F., Keene Beach, N., Camejo, P.Y., Oyserman, B.O., Kizaric, M., Khor, E.H., Noguera, D.R., McMahon, K.D., 2021. Metabolic differentiation of co-occurring Accumulibacter clades revealed through genome-resolved metatranscriptomics. mSystems 6 (4), e0047421.

McIlroy, S.J., Albertsen, M., Andresen, E.K., Saunders, A.M., Kristiansen, R., Stokholm-Bjerregaard, M., Nielsen, K.L., Nielsen, P.H., 2014. ‘*Candidatus Competibacter*’-lineage genomes retrieved from metagenomes reveal functional metabolic diversity. The ISME Journal 8 (3), 613–624.

Medini, D., Donati, C., Tettelin, H., Masignani, V., Rappuoli, R., 2005. The microbial pan-genome. Current Opinion in Genetics & Development 15 (6), 589–594.

Minh, B.Q., Schmidt, H.A., Chernomor, O., Schrempf, D., Woodhams, M.D., von Haeseler, A., Lanfear, R., 2020. IQ-TREE 2: new models and efficient methods for phylogenetic inference in the genomic era. Molecular iology and Evolution 37 (5), 1530–1534.

Monzon, V., Paysan-Lafosse, T., Wood, V., Bateman, A., 2022. Reciprocal best structure hits: using AlphaFold models to discover distant homologues. Bioinformatics Advances 2 (1), vbac072.

Nielsen, P.H., McIlroy, S.J., Albertsen, M., Nierychlo, M., 2019. Re-evaluating the microbiology of the enhanced biological phosphorus removal process. Current Opinion in Biotechnology 57, 111–118.

Oehmen, A., Lemos, P.C., Carvalho, G., Yuan, Z., Keller, J., Blackall, L.L., Reis, M.A.M., 2007. Advances in enhanced biological phosphorus removal: From micro to macro scale. Water Research 41 (11), 2271–2300.

Oehmen, A., Zeng, R.J., Yuan, Z., Keller, J., 2005. Anaerobic metabolism of propionate by polyphosphate-accumulating organisms in enhanced biological phosphorus removal systems. Biotechnology and Bioengineering 91 (1), 43–53.

Oyserman, B.O., Moya, F., Lawson, C.E., Garcia, A.L., Vogt, M., Heffernen, M., Noguera, D.R., McMahon, K.D., 2016a. Ancestral genome reconstruction identifies the evolutionary basis for trait acquisition in polyphosphate accumulating bacteria. The ISME Journal 10 (12), 2931–2945.

Oyserman, B.O., Noguera, D.R., del Rio, T.G., Tringe, S.G., McMahon, K.D., 2016b. Metatranscriptomic insights on gene expression and regulatory controls in *Candidatus* Accumulibacter phosphatis. The ISME Journal 10 (4), 810–822.

Páez-Watson, T., van Loosdrecht, M.C.M., Wahl, S.A., 2023. Predicting the impact of temperature on metabolic fluxes using resource allocation modelling: Application to polyphosphate accumulating organisms. Water Research 228, 119365.

Pál, C., Papp, B., Lercher, M.J., 2005. Adaptive evolution of bacterial metabolic networks by horizontal gene transfer. Nature Genetics 37 (12), 1372–1375.

Parks, D.H., Imelfort, M., Skennerton, C.T., Hugenholtz, P., Tyson, G.W., 2015. CheckM: assessing the quality of microbial genomes recovered from isolates, single cells, and metagenomes. Genome Res. 25 (7), 1043–1055.

Petriglieri, F., Singleton, C., Peces, M., Petersen, J.F., Nierychlo, M., Nielsen, P.H., 2021. “*Candidatus* Dechloromonas phosphoritropha” and “*Ca.* D. phosphorivorans”, novel polyphosphate accumulating organisms abundant in wastewater treatment systems. The ISME Journal 15 (12), 3605–3614.

Petriglieri, F., Singleton, C.M., Kondrotaite, Z., Dueholm, M.K.D., McDaniel, E.A., McMahon, K.D., Nielsen, P.H., 2022. Reevaluation of the phylogenetic diversity and global distribution of the genus *Candidatus* Accumulibacter. mSystems 7 (3), e00016–00022.

Pruitt, K.D., Tatusova, T., Maglott, D.R., 2006. NCBI reference sequences (RefSeq): a curated non-redundant sequence database of genomes, transcripts and proteins. Nucleic Acids Research 35 (suppl_1), D61–D65.

Qiu, G., Liu, X., Saw, N., Law, Y., Zuniga-Montanez, R., Thi, S.S., Ngoc Nguyen, T.Q., Nielsen, P.H., Williams, R.B.H., Wuertz, S., 2020. Metabolic traits of *Candidatus* Accumulibacter clade IIF strain SCELSE-1 using amino acids as carbon sources for enhanced biological phosphorus removal. Environmental Science & Technology 54 (4), 2448–2458.

Qiu, G., Zuniga-Montanez, R., Law, Y., Thi, S.S., Nguyen, T.Q.N., Eganathan, K., Liu, X., Nielsen, P.H., Williams, R.B.H., Wuertz, S., 2019. Polyphosphate-accumulating organisms in full-scale tropical wastewater treatment plants use diverse carbon sources. Water Research 149, 496–510.

Qiu, G., Law, Y., Zuniga-Montanez, R., Deng, X., Lu, Y., Roy, S., Thi, S.S., Hoon, H.Y., Nguyen, T.Q.N., Eganathan, K., Liu, X., Nielsen, P.H., Williams, R.B.H., Wuertz, S., 2022. Global warming readiness: feasibility of enhanced biological phosphorus removal at 35°C. Water Research 216, 118301.

Ravenhall, M., Škunca, N., Lassalle, F., Dessimoz, C., 2015. Inferring horizontal gene transfer. PLoS Computational Biology 11 (5), e1004095.

Roy, S., Guanglei, Q., Zuniga-Montanez, R., Williams, R.B.H., Wuertz, S., 2021. Recent advances in understanding the ecophysiology of enhanced biological phosphorus removal. Current Opinion in Biotechnology 67, 166–174.

Rubio-Rincón, F.J., Weissbrodt, D.G., Lopez-Vazquez, C.M., Welles, L., Abbas, B., Albertsen, M., Nielsen, P.H., van Loosdrecht, M.C.M., Brdjanovic, D., 2019. "*Candidatus* Accumulibacter delftensis": A clade IC novel polyphosphate-accumulating organism without denitrifying activity on nitrate. Water Research 161, 136–151.

Santos-Beneit, F., 2015. The Pho regulon: a huge regulatory network in bacteria. Frontiers in Microbiology 6, 402.

Saunders, A.M., Mabbett, A.N., McEwan, A.G., Blackall, L.L., 2007. Proton motive force generation from stored polymers for the uptake of acetate under anaerobic conditions. FEMS Microbiology Letters 274 (2), 245–251.

Seviour, R.J., Mino, T., Onuki, M., 2003. The microbiology of biological phosphorus removal in activated sludge systems. FEMS Microbiology Reviews 27 (1), 99–127.

Shang, Y., Wang, X., Chen, Z., Lyu, Z., Lin, Z., Zheng, J., Wu, Y., Deng, Q., Yu, Z., Zhang, Y., Qu, D., 2020. Staphylococcus aureus PhoU homologs regulate persister formation and virulence. Frontiers in Microbiology 11, 865.

Shen, W., Le, S., Li, Y., Hu, F., 2016. SeqKit: a cross-platform and ultrafast toolkit for FASTA/Q file manipulation. PLoS One 11 (10), e0163962.

Singleton, C.M., Petriglieri, F., Kristensen, J.M., Kirkegaard, R.H., Michaelsen, T.Y., Andersen, M.H., Kondrotaite, Z., Karst, S.M., Dueholm, M.S., Nielsen, P.H., Albertsen, M., 2021. Connecting structure to function with the recovery of over 1000 high-quality metagenome-assembled genomes from activated sludge using long-read sequencing. Nature Communications 12 (1), 2009.

Song, J.-M., Guan, Z., Hu, J., Guo, C., Yang, Z., Wang, S., Liu, D., Wang, B., Lu, S., Zhou, R., Xie, W.-Z., Cheng, Y., Zhang, Y., Liu, K., Yang, Q.-Y., Chen, L.-L., Guo, L., 2020. Eight high-quality genomes reveal pan-genome architecture and ecotype differentiation of Brassica napus. Nature Plants 6 (1), 34–45.

Srinivasan, V.N., Li, G., Wang, D., Tooker, N.B., Dai, Z., Onnis-Hayden, A., Bott, C., Dombrowski, P., Schauer, P., Pinto, A., Gu, A.Z., 2021. Oligotyping and metagenomics reveal distinct Candidatus Accumulibacter communities in side-stream versus conventional full-scale enhanced biological phosphorus removal (EBPR) systems. Water Research 206, 117725.

Tettelin, H., Masignani, V., Cieslewicz, M.J., Donati, C., Medini, D., Ward, N.L., Angiuoli, S.V., Crabtree, J., Jones, A.L., Durkin, A.S., DeBoy, R.T., Davidsen, T.M., Mora, M., Scarselli, M., Margarit y Ros, I., Peterson, J.D., Hauser, C.R., Sundaram, J.P., Nelson, W.C., Madupu, R., Brinkac, L.M., Dodson, R.J., Rosovitz, M.J., Sullivan, S.A., Daugherty, S.C., Haft, D.H., Selengut, J., Gwinn, M.L., Zhou, L., Zafar, N., Khouri, H., Radune, D., Dimitrov, G., Watkins, K., O’Connor, K.J.B., Smith, S., Utterback, T.R., White, O., Rubens, C.E., Grandi, G., Madoff, L.C., Kasper, D.L., Telford, J.L., Wessels, M.R., Rappuoli, R., Fraser, C.M., 2005. Genome analysis of multiple pathogenic isolates of *Streptococcus agalactiae*: Implications for the microbial pan-genome. Proceedings of the National Academy of Sciences 102 (39), 13950–13955.

Tian, Y., Chen, H., Chen, L., Deng, X., Hu, Z., Wang, C., Wei, C., Qiu, G., Wuertz, S., 2022. Glycine adversely affects enhanced biological phosphorus removal. Water Research 209, 117894.

Turcotte, M.M., Corrin, M.S.C., Johnson, M.T.J., 2012. Adaptive evolution in ecological communities. PLoS Biology 10 (5), e1001332.

Ulrich, L.E., Zhulin, I.B., 2010. The MiST2 database: a comprehensive genomics resource on microbial signal transduction. Nucleic Acids Research 38 (Database issue), D401–407.

Wang, L., Oehmen, A., Le, C., Liu, J., Zhou Y. Defluviicoccus vanus glycogen-accumulating organisms (DvGAOs) are less competitive than polyphosphate-accumulating organisms (PAOs) at high temperature. ACS EST Water 2021, 1, 319–327.

Wang, Z., Guo, F., Mao, Y., Xia, Y., Zhang, T., 2014. Metabolic characteristics of a glycogen-accumulating organism in *Defluviicoccus* cluster II revealed by comparative Genomics. Microbial Ecology 68 (4), 716–728.

Willsky, G.R., Malamy, M.H., 1980. Characterization of two genetically separable inorganic phosphate transport systems in Escherichia coli. Journal of Bacteriology 144 (1), 356–365.

Zaremba-Niedzwiedzka, K., Viklund, J., Zhao, W., Ast, J., Sczyrba, A., Woyke, T., McMahon, K., Bertilsson, S., Stepanauskas, R., Andersson, S.G., 2013. Single-cell genomics reveal low recombination frequencies in freshwater bacteria of the SAR11 clade. Genome biology 14 (11), 1–14.

Zhang, A.-N., Mao, Y., Wang, Y., Zhang, T., 2019. Mining traits for the enrichment and isolation of not-yet-cultured populations. Microbiome 7 (1), 96.

Zhang, C., Guisasola, A., Baeza, J.A., 2022. A review on the integration of mainstream P-recovery strategies with enhanced biological phosphorus removal. Water Research 212, 118102.

Zhang, C., Chen, X., Han, M., Li, X., Chang, H., Ren, N., Ho, S.-H., 2023. Revealing the role of microalgae-bacteria niche for boosting wastewater treatment and energy reclamation in response to temperature. Environmental Science and Ecotechnology 14, 100230.

Zhang, Z., Zhang, L., Zhang, G., Zhao, Z., Wang, H., Ju, F., 2023. Deduplication improves cost-efficiency and yields of de novo assembly and binning of shotgun metagenomes in microbiome research. Microbiology Spectrum 0 (0), e04282–04222.

Zhao, W., Bi, X., Peng, Y., Bai, M., 2022. Research advances of the phosphorus-accumulating organisms of *Candidatus* Accumulibacter, *Dechloromonas* and *Tetrasphaera*: Metabolic mechanisms, applications and influencing factors. Chemosphere 307, 135675.

Zhou, Y., Pijuan, M., Zeng, R.J., Yuan, Z., 2009. Involvement of the TCA cycle in the anaerobic metabolism of polyphosphate accumulating organisms (PAOs). Water Research 43 (5), 1330–1340.

